# Panax ginseng enhances the effect of metoprolol in chronic heart failure by inhibiting autophagy

**DOI:** 10.1101/2024.03.26.586847

**Authors:** Niu Zichang, Han Xiaoling, Liu Ting, Jin Qi, Ouyang Minghui, Mao Haoping

## Abstract

**Context:** Panax ginseng is often used as an adjuvant therapy for heart failure (HF) patients. Metoprolol is widely used in patients with HF. However, there is currently no report on the combined effects of ginseng and metoprolol in patients with HF.

**Objective:** The objective of this study is to investigate the combined effects of ginseng and metoprolol in patients with HF and the exact mechanisms.

**Materials and methods:** A mouse myocardial HF model was used. The serum levels of CK and CK-MB were determined using an automated biochemical analyzer. LDH and cTnT levels were determined using enzyme-linked immunosorbent assays. Autophagy of myocardial cells was evaluated using transmission electron microscopy, and changes in signal pathway proteins related to autophagy were analyzed using Western blot.

**Results:** The combination of ginseng and metoprolol can increase the survival rate of HF mice, improve heart function, reduce heart damage, and reduce serum CK, CK-MB, LDH, and cTnT levels. The combination of ginseng and metoprolol reduces autophagy in myocardial cells, reduces the levels of autophagy-related proteins (LC3, p62, Beclin1, and Atg5), and increases the p-PI3K/PI3K, p-Akt/Akt, and p-mTOR/mTOR ratios.

**Discussion and conclusion:** Ginseng enhances the anti-HF effect of metoprolol. Its mechanism of action may be related to inhibition of autophagy mediated by the PI3K/Akt/mTOR pathway.

## Introduction

Chinese herbal medicine has been used to prevent, treat, or cure diseases. Ginseng (*Panax* ginseng), the king of traditional Chinese medicine, is one of the most famous herbs and is often used as a therapeutic supplement or functional food to improve human health, including restoring physical vitality, enhancing immunity, and reducing cancer risk. Studies have shown that ginseng has various biological activities, such as immunomodulatory activity1, anti-inflammatory and antioxidant activity2, blood lipid- and blood sugar-regulating activity3,4, and cardiovascular activity5,6,7, and it is widely used for the treatment of heart failure (HF)8.

HF is a complex syndrome characterized by dysfunction of ventricular filling, ventricular ejection, or both. It is a global public health problem affecting more than 26 million people worldwide9. Data from the National Health and Nutrition Examination Survey show that the incidence and prevalence of HF are increasing10,11. It is becoming one of the most frequent causes of death. HF treatment strategies have seen major improvements. Multiple drugs have been shown to improve the quality of life, cardiac function, and survival in patients with HF. Commonly used drugs include beta-adrenergic receptor blockers (β-blockers), angiotensin-converting enzyme inhibitors (ACEIs), angiotensin receptor blockers (ARBs), angiotensin receptor–neprilysin inhibitors (ARNIs), mineralocorticoid receptor antagonists (MRAs), and sodium-glucose co-transporter 2 (SGLT2) inhibitors12. Among these drugs, β-blockers are essential and widely used in various heart diseases, especially in acute myocardial infarction (AMI) and HF with reduced ejection fraction (HFrEF). During the 1980s and 1990s, several large randomized controlled trials (RCTs) were conducted, which promoted the use of β-blockers, reducing mortality and morbidity rates in patients with chronic myocardial infarction and HFrEF. Beta-blockers reduce all-cause cardiovascular mortality, the occurrence of sudden cardiac death, and hospitalization for HF in patients with HFrEF13,14,15. Therefore, β-blockers should be prescribed to all patients with HFrEF that are not restricted or intolerant.

The clinical use of P. ginseng and β-blockers such as metoprolol (Met) is widespread. As interactions between traditional Chinese medicine and Western drugs may have beneficial or detrimental effects, combination therapy of *P. ginseng* and β-blockers should receive more attention. Therefore, we conducted the present study to investigate the effects of combined treatment with P. ginseng and Met, a commonly used β-blocker, in mice with HF and the underlying mechanisms.

## Materials and methods

### Drugs and reagents

Met tartrate tablets (H32025391) were obtained from MedChemExpress. (Shanghai, China). The clinical dosage was 50–100 mg twice daily. We converted a commonly used dosage of Met in clinical practice to a dose of 6.25 mg/kg in the experiment. P. ginseng crude drug herb was purchased from Beijing Tongrentang Pharmaceutical Co., Ltd (Beijing, china). The clinical dosage of P. ginseng is 3–9 g (for an adult weighing 70 kg) according to the Chinese Pharmacopoeia 2015. We converted the dosage of P. ginseng to a dose of 2.6 mg/kg in the experiment. Enzyme-linked immunosorbent assay (ELISA) kits for mouse cTnT were purchased from Elabscience (China). ELISA kits for the serum biochemical indicators LDH, CK, and CK-MB were purchased from Shenzhen Mindray Biomedical Electronic Co., Ltd (Shenzhen, China). Rabbit monoclonal anti-LC3, anti-Beclin1, anti-Atg5, anti-PI3K, anti-p-PI3K, anti-Akt, anti-p-Akt, anti-mTOR, and anti-p-mTOR were purchased from Abcam (Cambridge, UK). Rabbit monoclonal anti-p62 was purchased from CST (Boston, USA). Goat anti-rabbit IgG was purchased from Beijing Zhongshan Jinqiao Biotechnology Co., Ltd.

### Preparation of P. ginseng for animal experiments

Ginseng was prepared from *P.* ginseng. It was refluxed with 60% ethanol twice. Then it was concentrated and dissolved in distilled cold water to prepare a stock solution (1 g crude herb/mL). The contents of the ginseng extract were analyzed using UHPLC-PDA.

### Ultra performance liquid chromatography analysis

A Waters Ultra Performance Liquid Chromatography (UPLC) system coupled with a PDA detector was used to perform the quantitative analysis of the target compounds. All separation was performed on an ACQUITY UPLC BEH C18 column (2.1×100 mm, 1.7 μm). The flow rate was set as 0.3 mL/min. The column temperature was 40°C. The mobile phase comprised (A) aqueous formic acid (0.1%, v/v) and (B) methanol. The following gradient elution program was used: 30–52% B at 0–4 min, 52–57% B at 4–6 min, 57–65% B at 6–7 min, 65–69% B at 7–8 min, 69–73% B at 8–9 min, 73–75% B at 7–10 min, 75–80% B at 10–11 min, 80–90% B at 11–13 min, and 90–30% B at 13–15 min. The post-run-time was held at 5 min. The scan wavelength was set at 203 nm and the injection volume was 4 μL16. The P. ginseng extract (50 mg/mL) was centrifuged and filtered through a 0.22 μm membrane. Ginsenoside Re, Ginsenoside Rg1, Ginsenoside Rf1, and Ginsenoside Rb1 were dissolved in methanol to produce 1 mg/mL stock solutions, which were serially diluted to draw the calibration curve. All working solutions were stored at 4°C until use.

### Animals

Male 8-week-old C57BL/6J mice (20 ± 2 g) were purchased from Beijing Vital River Laboratory Animal Technology Co., Ltd. (Beijing, China, Certificate No.: SCXK Jing 2014-0004). They were housed under controlled temperature (23–26°C) and humidity (40–60%) conditions with a 12/12 h light/dark cycle. This study was conducted according to the recommendations in the Guidance Suggestions for the Care and Use of Laboratory Animals issued by the Ministry of Science and Technology of China. The Laboratory Animal Ethics Committee approved the protocol of Tianjin University of Traditional Chinese Medicine (Permit Number: TCM-LAEC2020086).

### Coronary artery ligation

Mice were anesthetized by inhalation of tribromoethanol. Then they were endotracheally intubated and maintained with ventilator-assisted breathing. The respiratory parameters were set so that the respiratory rate was 133 times/min, the tidal volume was 0.2 mL, and the total tidal volume per minute was 26 mL. Next, the chest was opened at the intercostal space between the third and fourth sternal ribs via a left thoracotomy. An 8-0 silk suture was used to ligate the proximal left anterior descending coronary artery under the left auricle. After successful ligation, the apex of the heart turned white. After suturing the thorax closure, the mice were released and placed on an electric blanket. The surgical procedures were identical in sham control mice, except the left anterior descending coronary artery was not tied. The mice were randomly divided into the following seven groups: sham, HF, Met (6.25 mg/kg), P. ginseng (1.3 g/kg), P. ginseng (2.6 g/kg), 6.25 mg/kg Met + 1.3 g/kg P. ginseng, and 6.25 mg/kg Met + 2.6 g/kg P. ginseng.

### Echocardiography

Left ventricular function was noninvasively assessed using an ultra-high-resolution small animal ultrasound imaging system (Vevo 2100, VisualSonics, Toronto, ON, Canada) equipped with a 30-MHz transducer. The following parameters indicating cardiac function were measured with M-mode: left ventricular ejection fraction (LVEF), left ventricular fractional shortening (FS), left ventricular internal diameter at diastole (LVID,d), and left ventricular internal diameter at systole (LVID,s).

### Histopathological examination

After being treated for 28 days, the mice were sacrificed under anesthesia. The heart was harvested, fixed in 4% paraformaldehyde solution for more than 48 h, and embedded in paraffin. Next, 5-µm-thick sections were cut from each segment and stained with hematoxylin–eosin (HE) and Masson trichrome. We used a microscope (Zeiss, China) to observe and image the sections.

### Measurement of biochemical parameters

The mice in each group were treated for 28 days. Then blood samples were collected, and serum was separated by centrifugation at 3500 rpm for 15 min. Serum levels of CK, CK-MB, and LDH were determined using an automatic biochemical analyzer (Mindray, China). Serum levels of cTnT were measured using an ELISA kit.

### Western blot analysis

The protein levels of LC3, p62, Beclin1, Atg5, PI3K, p-PI3K, Akt, p-Akt, mTOR, and p-mTOR were determined using Western blot. Cold lysis buffer was used to extract the total protein from the hearts. After centrifugation, the protein concentration in the supernatants was determined with a BCA Protein Assay Kit (Thermo, USA). Equal amounts of protein were separated using SDS-PAGE and transferred to polyvinylidene difluoride membranes, which were washed with Tris-buffered saline (TBS), blocked with TBS containing 5% skim milk or bovine serum albumin, incubated overnight at 4°C with primary antibody in TBST, washed, incubated with secondary antibody, and washed again. Finally, the protein bands were visualized using a chemiluminescence reagent, and signals were recorded using a chemiluminescence instrument (Bio-Rad Laboratories Inc., Hercules, CA, USA). The signal intensities were quantified using ImageJ.

### Statistical analysis

Experimental data are presented as the mean ± SEM. Statistical analysis was performed with a *t*-test for two groups or one-way ANOVA for multiple groups using SPSS 26.0. Values of *P* < 0.05 were considered to indicate a statistically significant difference.

## Results

### The contents of the P. ginseng extract

The contents of the P. ginseng extract were analyzed using UHPLC-PDA. Ginsenoside Re, Ginsenoside Rg1, Ginsenoside Rf1, and Ginsenoside Rb1 were detected and quantified. Representative chromatograms of the standards and P. ginseng extract are shown in **Figure 1**. The results showed that the P. ginseng extract contained 0.49 ± 0.01 mg/g Ginsenoside Re, 2.55 ± 0.03 mg/g Ginsenoside Rg1, 0.50 ± 0.03 mg/g Ginsenoside Rf1. and 2.70 ± 0.01 mg/g Ginsenoside Rb1.

**Figure 1.**
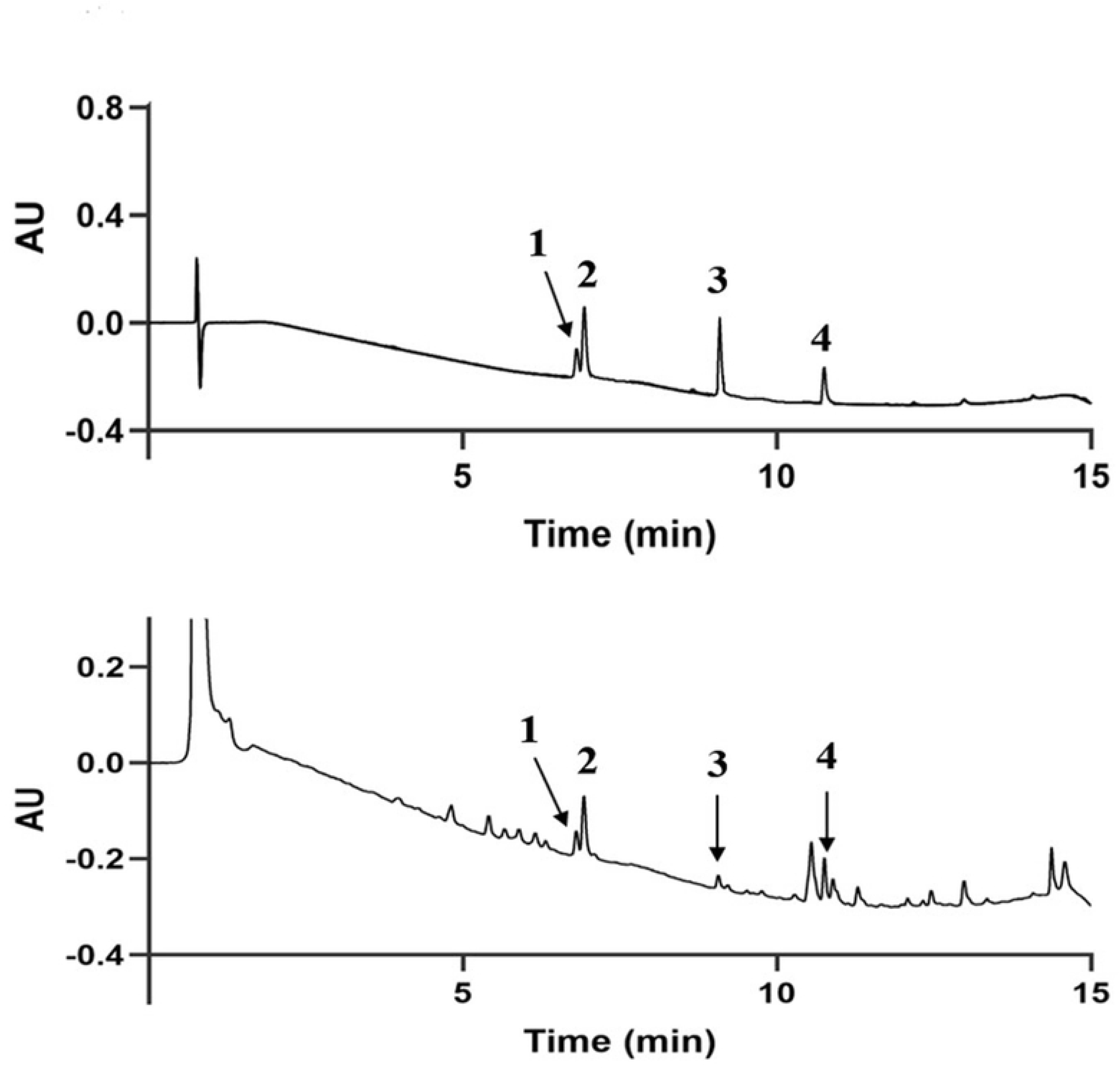
Typical chromatograms of (A) standard compounds and (B) samples. (1) Ginsenoside Re, (2) Ginsenoside Rg1, (3) Ginsenoside Rf1 and (4) Ginsenoside Rb1.

### P. ginseng increased survival in mice with HF

Our results showed that most deaths occurred within 7 days after surgery. Both P. ginseng and Met improved survival in mice. The survival rate was significantly higher in the 2.6 g/kg P. ginseng combined with Met group than in the Met group or the P. ginseng group at 28 days after surgery (**Figure 2**). This suggests that a combination of P. ginseng and Met would be beneficial for HF patients.

**Figure 2.**
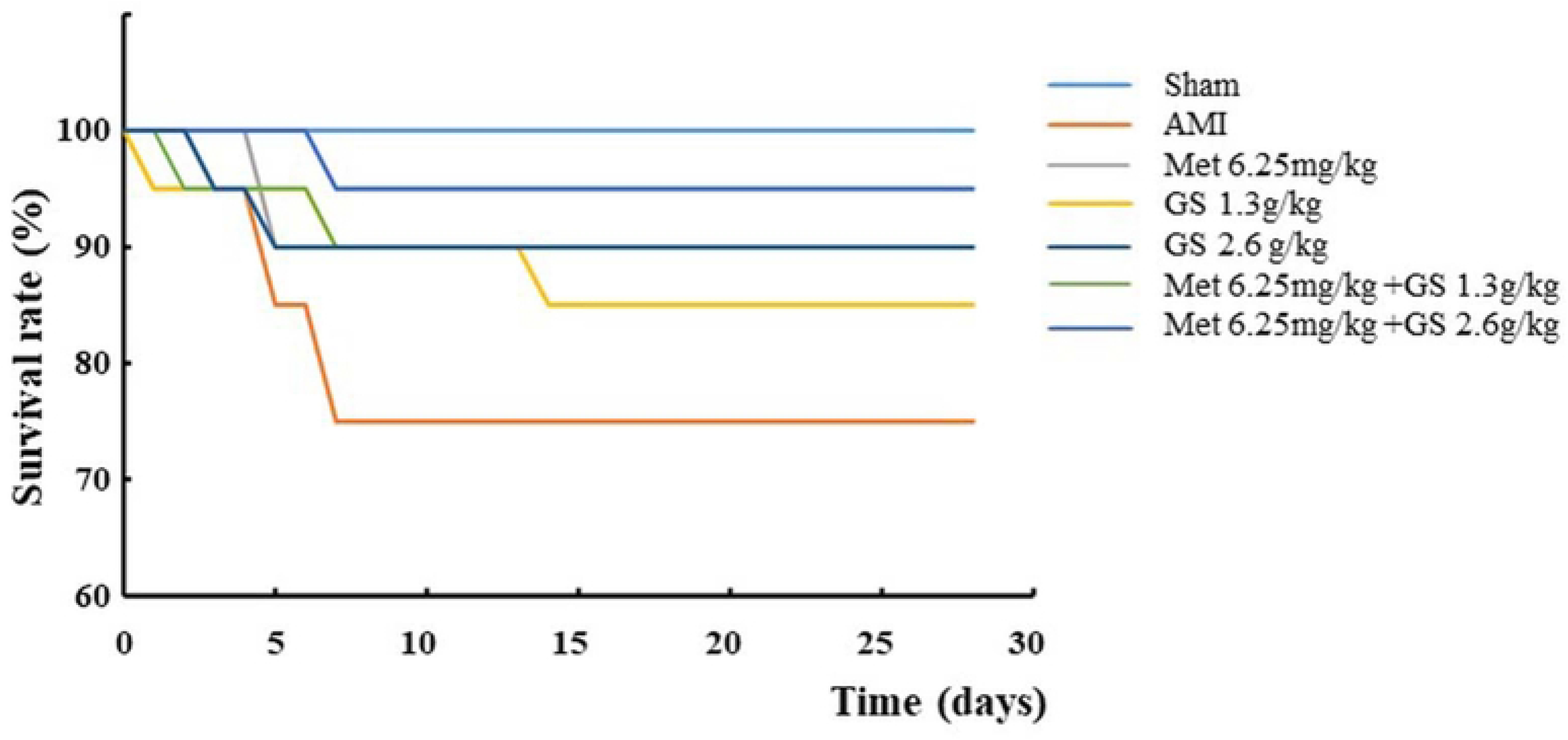
Effects of ginseng combined with Met on the survival rate of acute myocardial infarction-induced heart failure mice.

### P. ginseng combined with Met improved cardiac function

We subsequently employed echocardiography to assess cardiac function in HF mice (**Figure 3A**). Compared with the sham group, the EF and FS of the HF mice were significantly decreased (*P* < 0.01). A significant increase in EF was recorded when HF mice were treated with 6.25 mg/kg Met only. Mice treated with Met at a dose of 6.25 mg/kg also showed a significant increase in FS compared with HF mice (*P* < 0.05). Compared with HF mice, mice treated with P. ginseng at a dose of 2.6 g/kg showed a significant increase in EF and FS (*P* < 0.05 or *P* < 0.01). Furthermore, we found that the combination of 2.6 g/kg P. ginseng with Met significantly increased EF and FS compared with the Met and 2.6 g/kg P. ginseng groups (*P* < 0.05, **Figure 3B and C**). HF mice also showed increases in LVID,d and LVID,s (*P* < 0.01, **Figure 3D and E**), as well as increases in interventricular septum thickness in both diastole and systole (*P* < 0.05, **Figure 2F and G**). The combined use of P. ginseng and Met significantly reversed the changes in the above factors compared with the HF group (*P* < 0.01 or *P* < 0.05, **Figure 3D–G**). The above results indicate that *P.* ginseng combined with Met could promote the survival of HF-induced HF mice by improving cardiac function.

**Figure 3.**
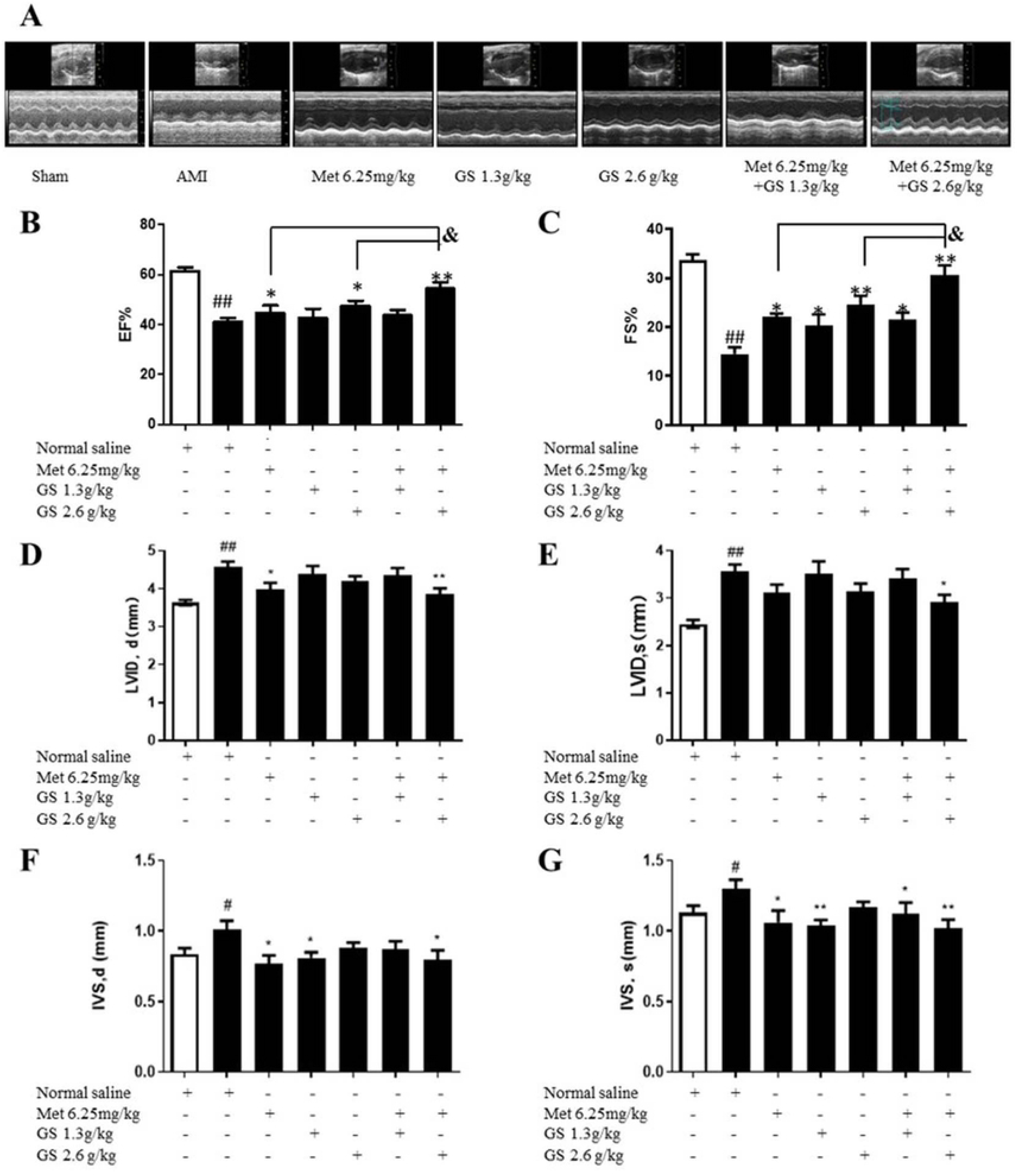
Typical echocardiograms (A), EF (B), FS (C), LVID,d (D), and LVID,s (E), IVS,d (F) and IVS.s (G) of heart failure mice treated with ginseng combined with Met. Data are represented as mean ± SEM (n = 7-10). #p < 0.05 or ##p < 0.01 vs. sham. *p < 0.05 or **p<0.01 vs. AMI. &p<0.05 vs. 2.6g/kg GS or 6.25mg/kg Met.

### P. ginseng combined with Met reduced heart damage and serum levels of CK, CK-MB, LDH, and cTnT in mice with HF

The HE staining results showed that the cardiomyocytes in the sham group were neatly arranged, the intercellular space was uniform, and the structure was complete. In the HF group, the myocardial cells were loosely arranged and disordered, the intercellular space was widened, and the myocardial stripes were broken and/or disappeared. Compared with the HF group, cardiomyopathy was significantly reduced in the 2.6 g/kg *P. ginseng* group and the 2.6 g/kg P. ginseng + 6.25 mg/kg Met group (**Figure 4A**). Masson staining showed that the myocardial tissue in the sham group was red and arranged orderly, and no collagen fibers appeared in the visual field. After AMI surgery, fibrosis and scarring appeared in the infarct border area of the myocardial tissue of the mice in each group, and the left ventricular wall became thinner, especially in the HF group. In addition, the remaining myocardial tissue in the HF group was less, the distribution of myocardial cells was uneven, and the structure was severely damaged. Compared with the HF group, the degree of myocardial fibrosis and the area of collagen fibers were significantly reduced in the treatment groups (**Figure 4B**).

**Figure 4.**
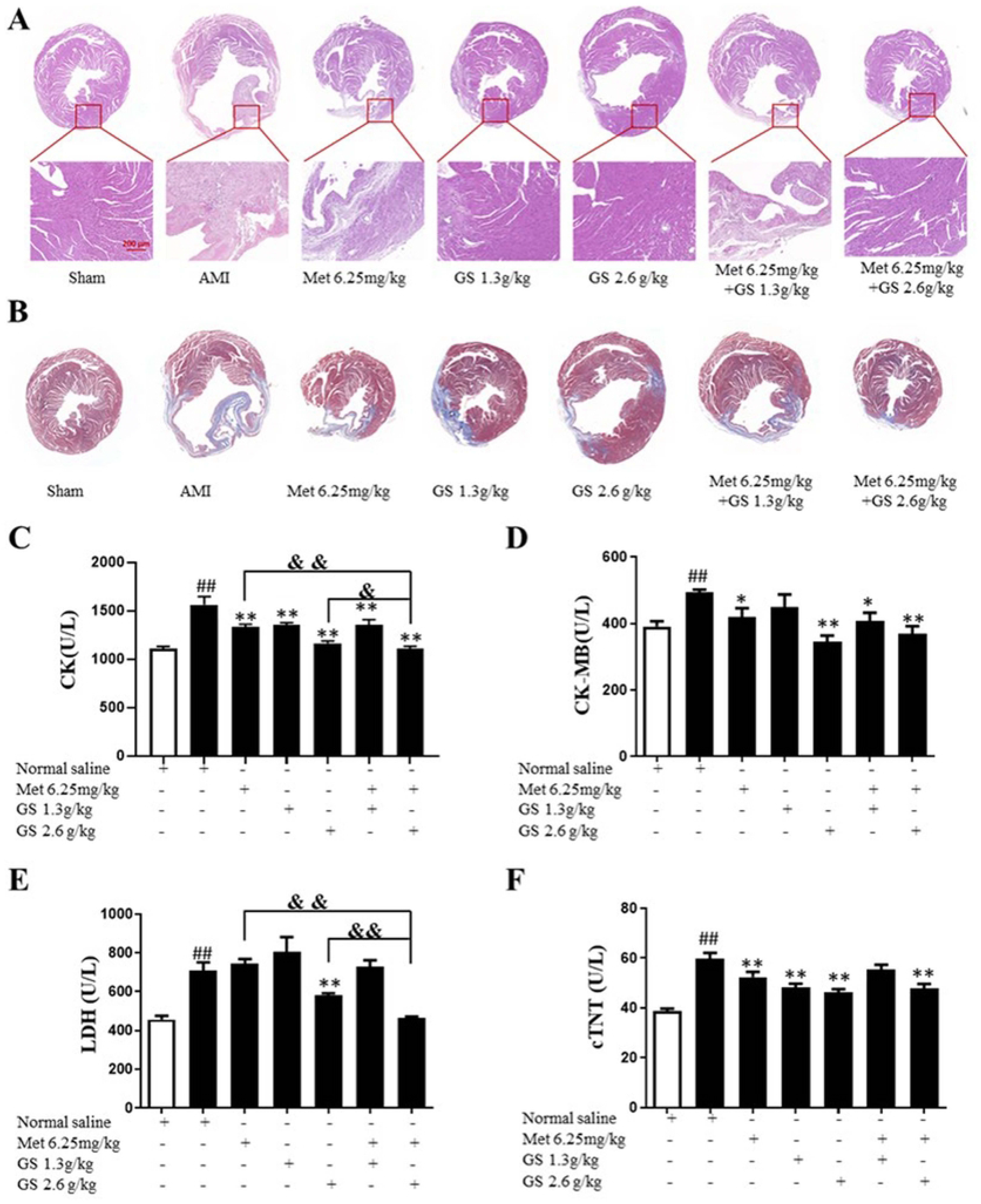
Effects of ginseng combined with Met on myocardial morphology and serum levels of CK, CK-MB, LDH, and cTnT in heart failure mice. (A) HE staining. (B) Masson staining. (C–F) Serum levels of CK (C), CK-MB (D), LDH (E), and cTnT (F). Data are represented as mean ± SEM (n = 7-10). ##p < 0.01 vs. sham. *p < 0.05 or **p<0.01 vs. AMI. &p<0.05 or &&<0.01 vs. 2.6g/kg GS or 6.25mg/kg Met.

An automatic biochemical analyzer was used to determine the serum levels of CK, CK-MB, and LDH on the 28th day after administration. As shown in Figure 3C, the serum CK content was significantly increased in the HF group compared with the sham group (P < 0.01). Compared with the HF group, the serum CK levels in the other groups were significantly decreased (P < 0.01). Compared with the 6.25 mg/kg Met group or the 2.6 g/kg P. ginseng group, the serum CK content in the 2.6 g/kg P. ginseng + 6.25 mg/kg Met group was significantly decreased (P < 0.01 or P < 0.05). As shown in **Figure 4D**, the serum CK-MB content in the HF group was significantly increased compared with the sham group (P < 0.01). Compared with the HF group, the serum levels of CK-MB were significantly reduced in the 6.25 mg/kg Met, 2.6 g/kg P. ginseng, 1.3 g/kg P. ginseng + 6.25 mg/kg Met, and 2.6 g/kg P. ginseng + 6.25 mg/kg Met groups (P < 0.05 or *P* < 0.01). As shown in **Figure 4E**, serum LDH levels were significantly increased in the HF group compared with the sham group (P < 0.01). Serum LDH levels were significantly decreased in the 2.6 g/kg P. ginseng + 6.25 mg/kg Met group compared with the 2.6 g/kg P. ginseng group or the 6.25 mg/kg Met group (P < 0.01). The serum cTnT level in the AMI group was significantly increased compared with the sham group (P < 0.01) (**Figure 4F**). The cTnT levels in the 6.25 mg/kg Met, 1.3 g/kg P. ginseng, 2.6 g/kg P. ginseng, and 2.6 g/kg P. ginseng + 6.25 mg/kg Met groups were significantly decreased (P < 0.01).

### Effects of P. ginseng combined with Met on autophagy-related proteins (LC3, p62, Beclin1, and Atg5) in mice with HF

Transmission electron microscopy revealed that AMI increased autophagosomes in cardiomyocytes and induced mitochondrial swelling and disorganization compared with sham-operated mice. Met reduced autophagosomes in cardiomyocytes and improved mitochondrial swelling and derangement compared with AMI mice. Both 1.3 g/kg *P. ginseng* and 2.6 g/kg P. ginseng reduced autophagosomes; however, no change in mitochondrial swelling and disorganization was observed compared with AMI mice. P. ginseng at 2.6 g/kg further reduced autophagosomes and improved mitochondrial swelling and derangement compared with Met alone (**Figure 5A**).

Further, we detected autophagy-related proteins. Compared with the sham group, the protein expression levels of LC3II/I, Beclin1, and Atg5 in the AMI group were significantly increased (P < 0.05 or *P* < 0.01). In contrast, the expression level of p62 was significantly decreased (P < 0.01). Compared with the AMI group, the protein expression levels of LC3bII/I, Beclin1, and Atg5 in the 2.6 g/kg P. ginseng + 6.25 mg/kg Met group were significantly decreased (P < 0.01); in contrast, the expression level of p62 was significantly increased (P < 0.05). Compared with the 6.25 mg/kg Met group, the protein expression levels of LC3bII/I, Beclin1, and Atg5 in the 2.6 g/kg P. ginseng + 6.25 mg/kg Met group were significantly decreased (P < 0.05 or *P* < 0.01); in contrast, the expression level of p62 was significantly increased (P < 0.05, **Figure 5B–F**).

**Figure 5.**
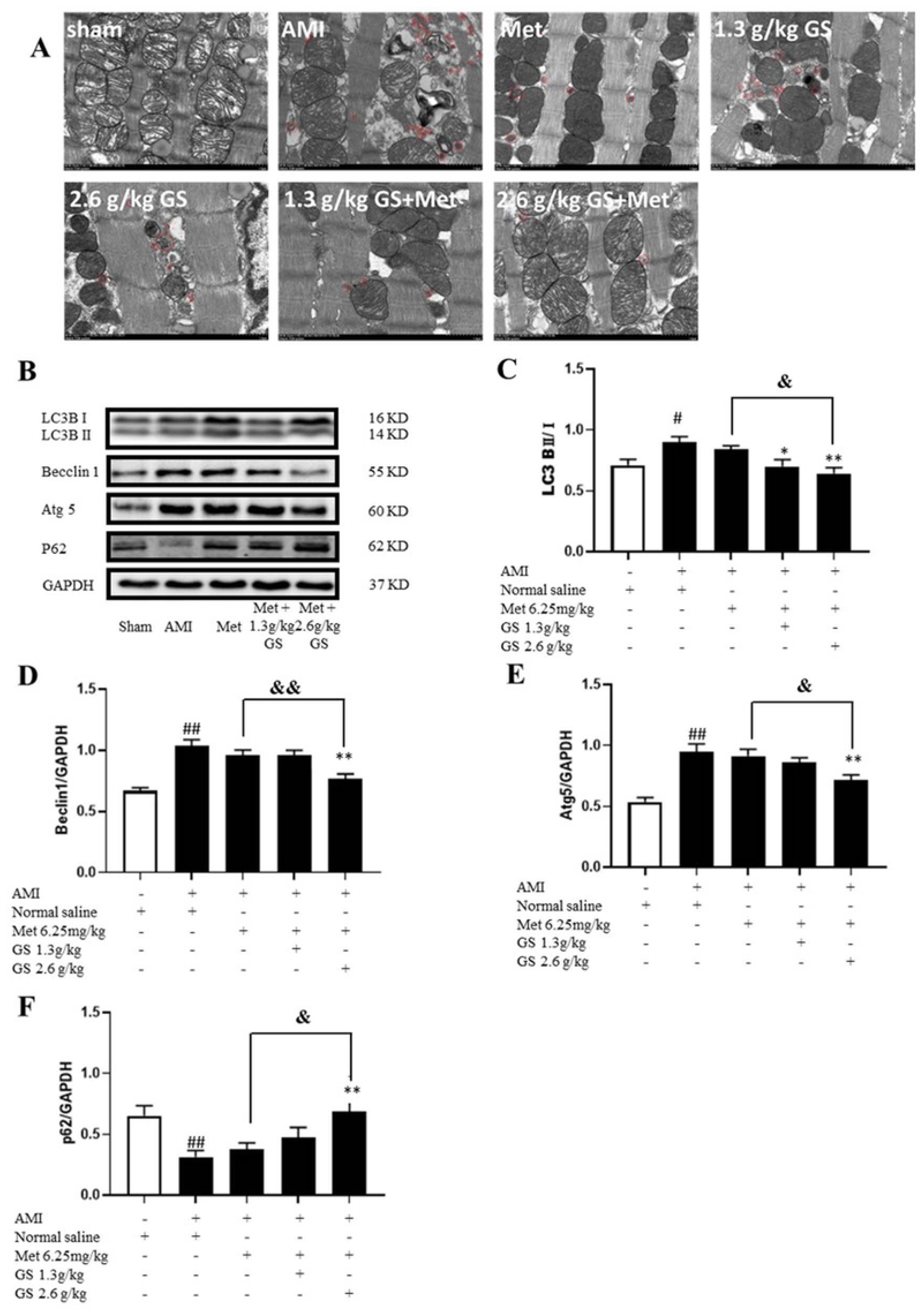
Effects of ginseng combined with Met on autophagy-related proteins (LC3BII/I, p62, Beclin1, and Atg5) in heart failure mice. (A) Typical transmission electron microscopy image. (B–F) Western blot analysis of autophagy-related proteins (LC3BII/I, p62, Beclin1, and Atg5). Data are represented as mean ± SEM (n = 7-10). #p < 0.05 or ##p < 0.01 vs. sham. *p < 0.05 or **p<0.01 vs. AMI. &p<0.05 or &&<0.01 vs. 2.6g/kg GS or 6.25mg/kg Met.

### Effects of P. ginseng combined with Met on PI3K, p-PI3K, p-Akt, Akt, p-mTOR, and mTOR in the upstream signaling pathway of autophagy in mice with HF

Western blot results for p-PI3K/PI3K, p-Akt/Akt, and p-mTOR/mTOR are shown in **Figure 6A**. The ratios of p-PI3K/PI3K, p-Akt/Akt, and p-mTOR/mTOR were significantly decreased in the AMI group compared with the sham group (P < 0.01), and they were significantly higher in the 2.6 g/kg P. ginseng + 6.25 mg/kg Met group than in the AMI group (P < 0.01). There were no significant differences between the other groups and the AMI group. The p-Akt/Akt and p-mTOR/mTOR ratios were significantly higher in the 2.6 g/kg P. ginseng + 6.25 mg/kg Met group than in the Met group (P < 0.01) (**Figure 6B**), while the p-PI3K/PI3K ratio was not significantly different (**Figure 6C and D**).

**Figure 6.**
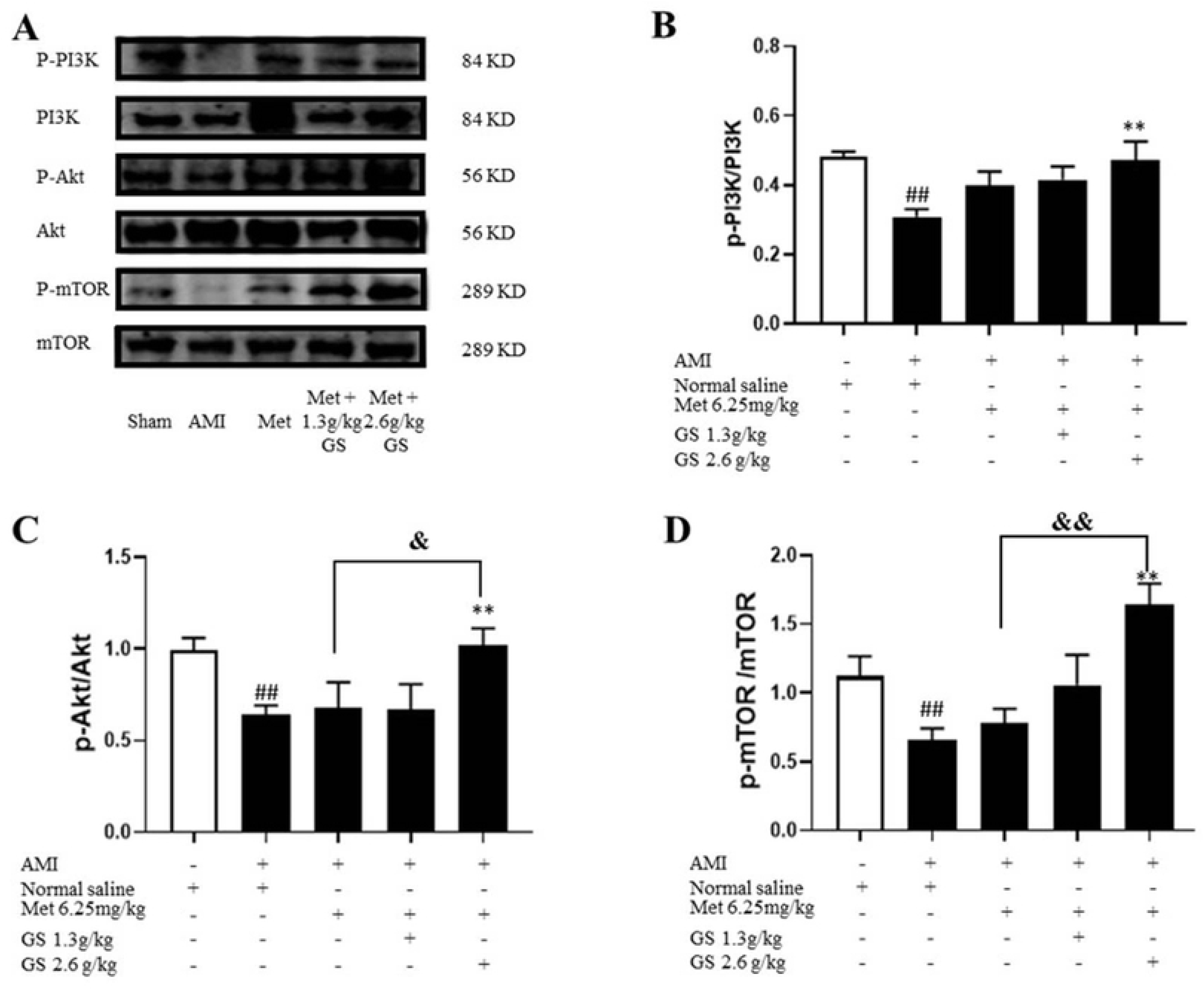
Effects of ginseng combined with Met on PI3K, p-PI3K, Akt, p-Akt, mTOR, and p-mTOR levels in heart failure mice. Data are represented as mean ± SEM (n = 7-10). ##p < 0.01 vs. sham. *p<0.05 or **p<0.01 vs. HF. &p<0.05 or &&<0.01 vs. 2.6g/kg GS or 6.25mg/kg Met.

## Discussion

HF has been reported to affect approximately 6.5 million US adults, with approximately 1 million hospitalizations annually, approximately 50% of which are caused by HFrEF17. One in six patients with HFrEF has a high risk of hospitalization and death due to HF recurrence within 2 years18. Therefore, the treatment of HFrEF plays an essential role in human health.

Modern medicine has seen major improvements. New therapies and drugs such as MRAs and SGLT2 inhibitors have been developed to improve clinical outcomes in patients with HFrEF. Beta-blockers are used in the treatment of cardiovascular diseases and remain essential in HFrEF treatment. In a review, Philip et al. summarized the current evidence underlying the use of β-blockers in the treatment of HF and found that the current evidence strongly supports the use of β-blockers in the treatment of HFrEF19. Current HFrEF clinical practice guidelines recommend the combined use of these medications. Combinations of ARNIs, β-blockers, MRAs, and SGLT2 inhibitors have been reported to reduce the occurrence of hospitalization and cardiovascular death due to HF, as well as all-cause mortality20. Therefore, β-blockers are widely used in the treatment of HFrEF. Beta-blockers reduce myocardial oxygen consumption by reducing the heart rate, blood pressure, and myocardial contractility21. Therefore, β-blockers are considered the cornerstone of HF treatment. Met is a conventional β-blocker used in HF treatment, although several new β-blockers with different clinical features have emerged. Met is widely reported to be effective in HF and is associated with lower HF readmission and mortality rates in hospitalized patients or within 1 year of HF onset22-23. Met is a commonly used selective β1-adrenergic receptor blocker. Intravenous administration of Met has been reported to reduce infarct size and improve long-term cardiac function after AMI in clinical trials24-25 and experimental studies26.

Autophagy is an essential process involved in HF pathology. Autophagy is reported to have double-edged sword-like effects in myocardial tissue; moderate autophagy is necessary for cardiomyocytes to survive and maintain the heart’s normal function, whereas excessive activation of autophagy may result in decreased cardiac function and even HF27. Several signaling pathways regulating autophagy have been identified, including the mTOR28, AMPK29, and ROS signaling pathways30. The mTOR pathway plays a significant role in HF. Inhibition of mTOR-related autophagy has been shown to have the potential for HF treatment and prevention28,31. To further elucidate the mechanisms by which P. ginseng combined with Met improves cardiac function, we examined the protein expression levels of p-PI3K/PI3K, p-Akt/Akt, and p-mTOR/mTOR. Our results demonstrate that P. ginseng combined with Met significantly inhibited PI3K, Akt, and mTOR phosphorylation.

The use of β-blockers which counteract neurohormonal hyperactivation is an established cornerstone for the treatment of HF. Thus, β-blockers are recommended as first-line therapy in the presence of left ventricular dysfunction and must be titrated to the maximum tolerated dose to achieve their full benefits. Met used once daily in addition to optimum standard therapy was reported by many randomized trials to effectively reduce all-cause mortality and the occurrence of sudden death compared with the placebo group32.

HF is one of the most prevalent cardiovascular diseases in the 21st century, with high morbidity and mortality rates. Myocardial ischemia and infarction caused by coronary heart disease are the most common causes of HF. Advances in medical technology have enabled more patients with acute cardiovascular disease to survive and develop post-myocardial infarction HF, which eventually develops into chronic HF. Early revascularization can minimize myocardial ischemic necrosis in patients with AMI, thereby reducing the occurrence or severity of HF. At the same time, early standardized drug treatment, including β-blockers, ACEIs, and ARBs, can further reduce the occurrence of HF. Among them, β-blockers are recommended to be used in the early stages of AMI at low doses due to their positive effects, reducing myocardial infarction size, recurrent myocardial ischemia, and the risk of reinfarction33. The updated guideline for the management of HF released in 2017 showed that the dosages of Met succinate were 12.5–25 mg QD for initial usage and 200 mg QD for maximum usage, and the mean dose administered in clinical trials was 159 mg QD34. However, the dosing of β-blockers such as Met in the clinic varied markedly, often falling short of guideline recommendations. A national assessment of early β-blocker therapy in patients with AMI in China’s PEACE study reported substantial underuse of early β-blocker therapy in ideal candidates with AMI from 2001 to 201135. A study published in 2021 also reported that the early use of oral β-blockers in acute coronary syndrome patients was generally insufficient, with huge differences among different hospitals in China36. In another study, it was reported that 88% of Met users did not take the recommended daily dose (200 mg)37. Therefore, the effects of P. ginseng combined with different Met doses are worth studying. In the present study, we used 6.25 mg/kg Met to investigate the effects of the combination of P. ginseng and Met in AMI-induced HF mice. In our preliminary experiment, 6.25 mg/kg Met showed no significant effect on HF after AMI, so we used 6.25 mg/kg Met with P. ginseng first to investigate the effect of P. ginseng combined with a low dosage of Met since Met is often underused. To the best of our knowledge, this is the first report of the effect of the combination of P. ginseng and a low dose of Met in HF after AMI. Our results demonstrate that co-treatment with P. ginseng and Met could be beneficial.

## Conclusion

We demonstrated that a combination of P. ginseng with 6.25 mg/kg Met increased the survival rate and heart function in AMI-induced HF mice by inhibiting autophagy through the PI3K/Akt/mTOR pathway.

## Acknowledgments

We thank Key laboratory of Pharmacology of Traditional Chinese Medical Formulea for its linguistic assistance during our work.

## Consent for publication

All authors agree to publish this article.

